# Blood donor variability is a modulatory factor for *P. falciparum* invasion phenotyping assays

**DOI:** 10.1101/2020.05.27.116939

**Authors:** Laty G. Thiam, Prince B. Nyarko, Kwadwo A. Kusi, Makhtar Niang, Yaw Aniweh, Gordon A. Awandare

## Abstract

Human erythrocytes are indispensable for *Plasmodium falciparum* development. Unlike other eukaryotic cells, there is no existing erythroid cell line capable of supporting long-term *P. falciparum in vitro* experiments. Consequently, invasion phenotyping experiments rely on erythrocytes of different backgrounds. However, the contribution of the erythrocytes variation in influencing invasion rates remains unknown, which presents a challenge for conducting large-scale comparative studies. Here, we used erythrocytes of different blood groups harboring different hemoglobin genotypes to assess the relative contribution of blood donor variability in *P. falciparum* invasion phenotyping assays. For each donor, we investigated the relationship between parasite invasion phenotypes and erythrocyte phenotypic characteristics, including; the expression levels of surface receptors (e.g. the human glycophorins A and C, the complement receptor 1 and decay accelerating factor), blood groups (e.g. ABO/Rh system), and hemoglobin genotypes (e.g. AA, AS and AC). Across all donors, there were significant differences in invasion efficiency following treatment with either neuraminidase, trypsin or chymotrypsin relative to the control erythrocytes. Primarily, we showed that the levels of key erythrocyte surface receptors and their sensitivity to enzyme treatment, significantly differed across donors. However, invasion efficiency correlated neither with susceptibility to enzyme treatment nor with the levels of the selected erythrocyte surface receptors. Upon further analysis, we found no relationship between *P. falciparum* invasion phenotype and blood group or hemoglobin genotype.

**Importance:** Assays to decipher *P. falciparum* invasion phenotypes are of great importance in the quest for an efficient malaria vaccine. Malaria associated mortality is mainly attributed to the blood stage of the parasite’s life cycle, a major focus of vaccine development strategies. Further, testing and validating blood stage vaccines necessitates conducting large-scale studies in endemic countries. However, comparing results from such studies is challenged by the lack of standard assays. As human erythrocytes play a pivotal role in *P. falciparum* invasion assays, the need to investigate the effect of blood donor variability in the outcome of such assays is apparent. The significance of our study is in reporting the absence of relationship between *P. falciparum* invasion efficiency and commonly shared erythrocyte features across different erythrocyte donors, therefore emphasizing the need to consider erythrocyte donor uniformity and to anticipate challenges associated to blood donor variability in early stages of large-scale study design.

## Introduction

Malaria continues to be a global public health burden, causing over two hundred million cases annually and accounting for hundreds of thousands of deaths every year (1). Alone, *Plasmodium falciparum* accounts for more than 90% of the malaria-related mortality globally, primarily occurring in children and pregnant women living in sub-Saharan Africa (2). Malaria-associated pathologies only manifest during the blood stage of the parasite’s life cycle. This stage is characterized by repeated rounds of asexual replications within the host erythrocyte, following the parasite’s egress from the hepatocytes. *P. falciparum* merozoites have the sole purpose to invade erythrocytes and perpetuate the asexual multiplication (3). Given their importance in the parasite’s successful invasion and further multiplication within the host cell, merozoite antigens, and particularly invasion-related antigens present attractive blood-stage vaccine targets. Thus, unravelling the nature of ligand-receptor interactions involved in erythrocyte invasion is essential for malaria vaccine development.

Although recent studies have enabled considerable progress in our understanding of the molecular basis of erythrocyte invasion by *Plasmodium* parasites (4, 5), little is known about the actual contribution of the host cell. Although there is clarity on the redundancy of ligand-receptor interactions involved in invasion, the functional relevance of some of these interactions are uncertain. Such interactions are presumed to be involved in signal transduction on either side between the parasite and the host erythrocyte (6, 7). However, pioneering reports on the major invasion profiles of *P. falciparum* clinical isolates across various malaria endemic countries have led to the hypothesis that *P. falciparum* invasion profiles are driven by the intensity of ongoing transmission in any given area (8–16). This proposition has been challenged by recent findings, which have shown no relationship between endemicity and invasion profile when parasites from countries of varying endemicity were subjected to similar protocols (16). This emphasizes that conducting large-scale *P. falciparum* phenotyping studies may inevitably require standardized protocols to allow comparisons across sites. One of the major drawbacks that may preclude the design of such assays is the lack of consistency in the usage of donor erythrocytes (17). Human erythrocyte polymorphisms have been shown to be associated with the distribution of *P. falciparum* globally (18, 19). This heterogeneity may account for the differences in the reported invasion profiles using erythrocytes of different origins.

In addition, despite the progress made in generating immortalized erythroid cell-lines retaining a mature phenotype, upscaling the production of these cells for universal usage is challenging (20–25). It is therefore of utmost importance to investigate the contribution of variation in donor erythrocytes in characterizing *P. falciparum* phenotypic diversity. Here, we present results from investigations aimed at assessing the relative contribution of blood donor variability in *P. falciparum* invasion phenotyping assays (IPAs).

We showed that, following enzyme treatments, there was a significant variation in the parasites’ invasion efficiency into individual donor erythrocytes. Moreover, the levels of key erythrocyte surface receptors and their sensitivity to enzyme treatment significantly differed across donors. However, invasion efficiency did not significantly correlate with susceptibility to enzyme treatment or the expression levels of the selected erythrocyte surface receptors.

## Materials and Methods

### Demographic and hematological characteristics of the study participants

The use of human erythrocytes for this study was approved by the Institutional Review Board (IRB) of the Noguchi Memorial Institute for Medical Research Ethics Committee, University of Ghana (IRB00001276) and the Ghana Health Services (GHS) ethical review committee (GHC-ERC:005/12/2017). Written informed consent was obtained from all participants. Blood samples were collected from twenty non-related asymptomatic adults, comprising fifteen males and five females, all resident in Ghana. Donors were questioned about their most recent clinically diagnosed malaria symptoms and to eliminate possible confounders, only individuals with no recent history of clinical malaria (at least two years) were considered. Additionally, erythrocytes from a single donor, used for routine parasite culturing, were included in all assays to normalize the resulting parasitemia. All but one sample were subjected to clinical diagnosis to screen for possible hemoglobin disorders while all samples were typed for ABO/Rh blood group presented in Table A1. In brief, the majority of the donors (14/20) presented a normal hemoglobin genotype (AA), while four donors had sickle cell trait (AS) and two other donors had an AC genotype. Blood group O^+^ was the commonest in all donors (10/20), followed by the A^+^ and B^+^ (5 and 4, respectively), while O^−^ was the least common blood group in the study participants. Of all donors, only one presented a severe deficiency of the G6PD expression. Full blood count was also performed to assess other hematological indices (Table A1). All samples were collected in ACD vacutainers (BD Biosciences, USA) and washed three times with RPMI 1640 (Sigma Aldrich, UK) to separate the erythrocytes from the other blood components. For each sample, part of the resulting erythrocyte pellet was used for IPAs while the remaining was cryopreserved for further experiments.

### *P. falciparum* isolates and culture conditions

All *P. falciparum* strains were maintained in culture at 4% hematocrit in complete parasite medium (RPMI 1640 containing 25 mM HEPES, 0.5% Albumax II, 2 mg/mL sodium bicarbonate and 50 μg/mL Gentamicin) and incubated at 37°C in an atmosphere of 5.5% CO_2_, 2% O_2_ and balance N_2_ gas mixture. Parasites were maintained in culture using a single donor O^+^ erythrocytes and routinely synchronized using 5% D-Sorbitol (Sigma Aldrich, UK).

### Invasion assay set up

For each donor, erythrocytes were either untreated or treated with different enzymes, including neuraminidase (250 mU/mL), trypsin (1 mg/mL) or chymotrypsin (1mg/mL); and labelled with 20 μM carboxyfluorescein diacetate succinimidyl ester (CFDA-SE) as described earlier (26). For each isolate, schizont stage parasites were inoculated at a 1:1 ratio into fresh enzyme-treated and labelled erythrocytes from a given donor. Experiments were conducted in triplicates in 96 well plates and repeated at least two times. Parasites were incubated for about 24 hours, after which the cells were stained with 5 μM Hoechst 33342 to label the parasite’s DNA and the invasion efficiency was assessed by flow cytometry. The percentage of erythrocytes positive for both dyes was recorded as the invasion efficiency and the parasite's invasion phenotype was determined by comparing invasion rates in enzyme-treated erythrocytes to that of untreated cells. To minimize the effect of any possible confounders that may arise during the sample processing or assay set up, erythrocytes from a single donor, used for routine parasite culturing, were included in all assays and used to finally normalize the resulting parasitemia.

### Characterization of erythrocyte surface receptors

The surface expression of selected erythrocyte receptors known or predicted to be involved in invasion was quantified by flow cytometry using specific monoclonal antibodies. Freshly washed erythrocytes were diluted to 1% hematocrit in 1X PBS containing 1% BSA and coincubated for an hour with antibodies against the human glycophorin AB (GYPAB) (1:400, Sigma Aldrich UK), GYPA (Clone E4, 1:100, Santa Cruz Biotech, USA), GYPC (Clone E3, 1:100, Santa Cruz Biotechnology, USA), complement receptor 1 (CR1, Clone J3D3, 1:50, Santa Cruz Biotechnology, USA) or Decay-accelerating factor (DAF, Clone NaM16-4D3, 1:50, Santa Cruz Biotechnology, USA). The erythrocyte pellets were collected by centrifugation at 2000 rpm, washed twice with 1X PBS and subsequently coincubated with anti-mouse antibodies conjugated with either Alexa Fluor 700, APC or PE (Santa Cruz Biotechnology, USA). All incubations were done at 37°C for an hour and protected from light exposure. The data were acquired using a BD LSR Fortessa X-20 flow cytometer (BD Biosciences, Belgium) and analyzed using FlowJo v10.5.0 (FlowJo, LLC, Ashland OR) and GraphPad Prism v.8.01 (GraphPad Software Inc., La Jolla, CA, USA).

### Antibody-dependent invasion inhibition assays

To ascertain the relative contribution of receptor density in the invasion efficiency, CFDA-labelled erythrocytes from different donors were pre-incubated at 37°C for an hour with different concentrations of antibodies against the human GYPC, CR1 or DAF. Antibody dilutions (range 0.3125 to 5 μg/mL) were optimized to prevent formation of cell aggregates. The erythrocytes were pelleted by centrifugation at 2000 rpm for 3 minutes, and washed with PBS prior to the addition of schizont-infected cells. The antibody-bound erythrocytes were then coincubated at 2% hematocrit with equal volumes of parasitized erythrocytes under normal culture conditions. The parasites’ invasion rates were assessed by flow cytometry upon reinvasion.

### Statistical analyses

All statistics were performed using GraphPad Prism v.8.01 (GraphPad Software, Inc.). The data were analyzed as the mean and standard error of pooled data from at least two independent experiments conducted each in triplicates. Statistical significance of invasion efficiency into enzyme-treated erythrocytes from different donors was ascertained using the Two-way ANOVA, coupled with the Tukey's multiple comparison test for pairwise analysis. Spearman correlation test was used to ascertain the relationship between variables while the Kruskal Wallis and Mann-Whitney tests were used to compare different groups.

## Results

### Invasion phenotypes of *P. falciparum* in erythrocytes from different donors

To investigate the effects of blood donor variability on *P. falciparum* invasion phenotype, we determined the invasion efficiency of various *P. falciparum* isolates in erythrocytes from different donors. First, we determined the sensitivities of the different donor erythrocytes to neuraminidase, which removes sialic acid (SA) residues on glycophorins, and trypsin and chymotrypsin which selectively cleave peptide backbones of other receptors (27, 28). Measurement of receptor surface expression levels showed a range of sensitivities to the three enzymes across donors (Fig 1). We conducted assays to determine the invasion of three newly culture-adapted isolates (MISA010, MISA011 and MISA018) into enzyme-treated erythrocytes from twenty donors. Laboratory strains of *P. falciparum* (3D7, Dd2 and W2mef), with known invasion phenotypes, were also assessed. Individually, all parasite strains tested showed differences in invasion efficiency across donors (Tables A2–A7), and these variations remained after the data were normalized and pooled for each donor (Fig 2).

**Figure 1:**
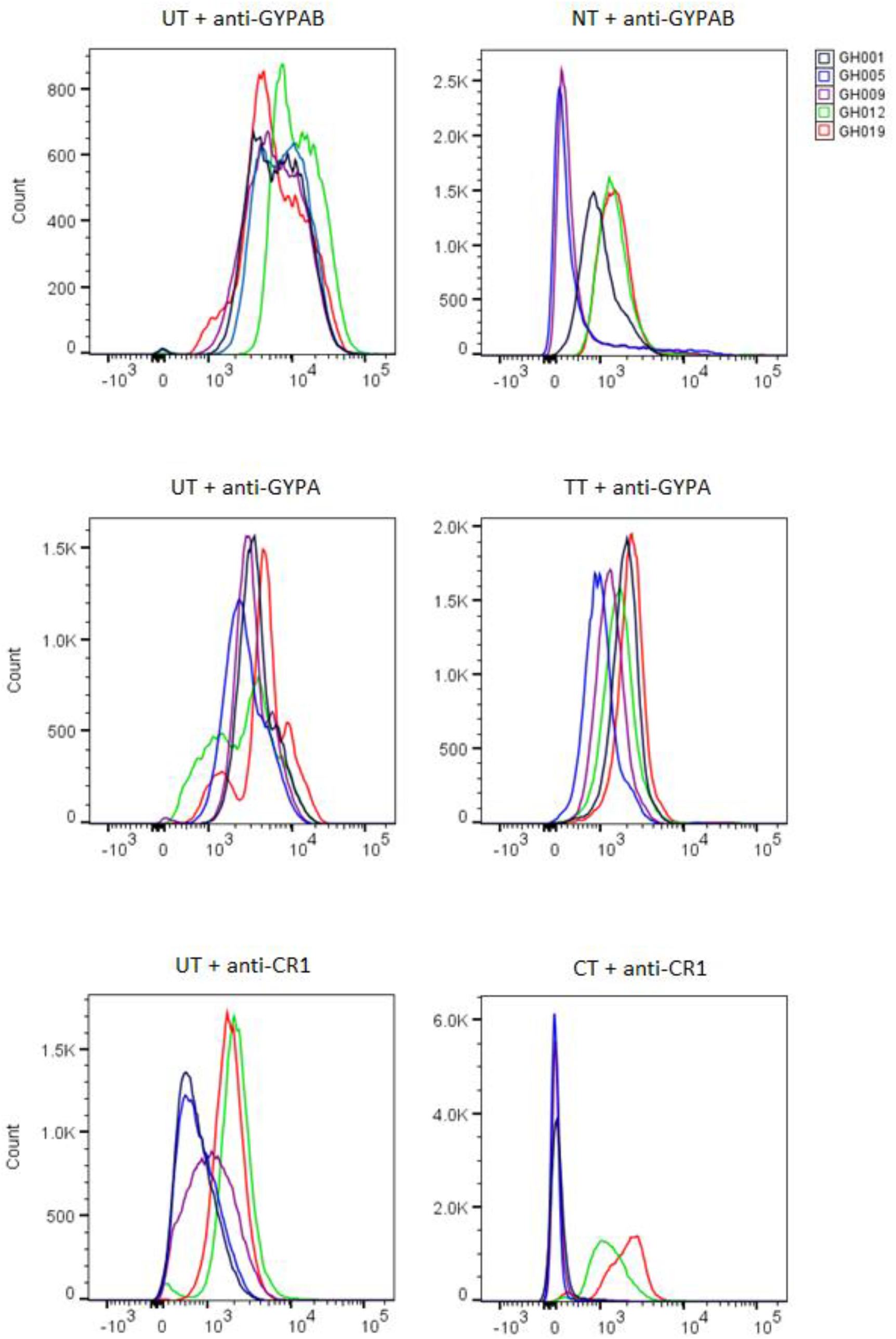
Efficacy of enzyme treatment of erythrocytes from different donors. Histogram plots showing the fluorescent intensity associated with antibodies against specific erythrocyte receptors prior to (left panel), and after (right panel) treatment with different enzymes (NT: neuraminidase treatment, TT: trypsin treatment, CT: chymotrypsin treatment). Target erythrocytes from five different donors were coincubated for an hour with mouse anti-human GYPAB (top panel), GYPA (middle panel) or CR1 (bottom panel), then washed twice and incubated with goat anti-mouse-PE conjugated antibodies. The data was collected with a BD LSR Fortessa X-20 flow cytometer and graphs plotted using FlowJo v.10.01.

**Figure 2:**
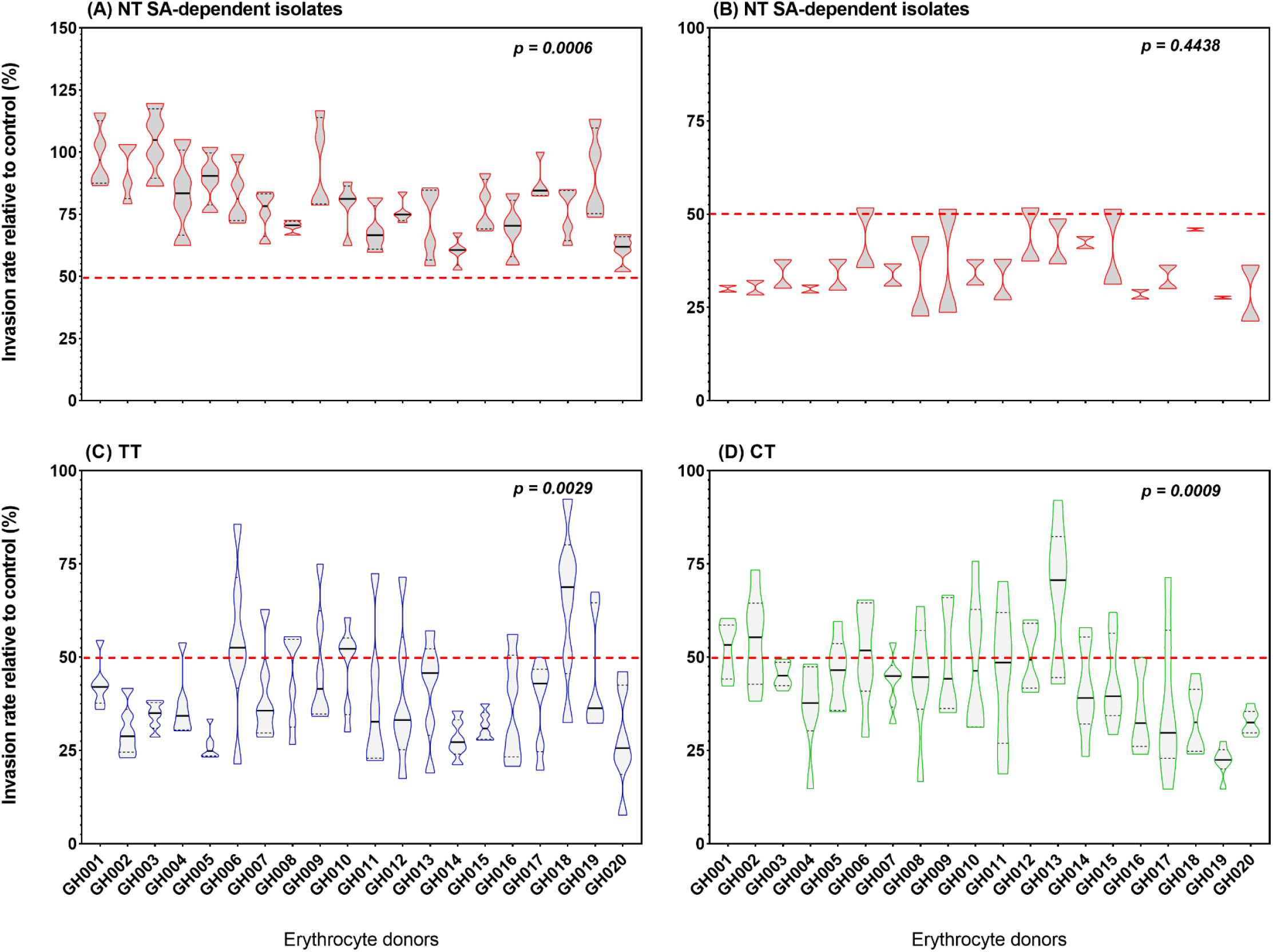
*P. falciparum* invasion phenotypes into erythrocytes from twenty different donors. The assay was set up along with a single donor erythrocyte used as a control for normalizing the resulting parasitemia. Presented are the means and standard errors of the invasion efficiencies from six different *P. falciparum* isolates assayed in triplicates in two independent experiments. The top panel graphs represent the invasion phenotypes of both SA-dependent (A) and independent (B) following the neuraminidase treatment (NT) of donor erythrocytes. The bottom panel graphs and (D) represent the invasion phenotypes obtained following trypsin (TT) and chymotrypsin (CT) treatment respectively. The red-dotted line indicates the sensitivity cut off for each given treatment. Statistical tests were performed using the Friedman test followed by Tukey’s multiple comparison test.

For neuraminidase-treated erythrocytes, while no significant difference was observed between SA-dependent parasites (Fig 2A, *P = 0.4438*), invasion rates significantly differed between the SA-independent isolates (Fig 2B, *P = 0.0006*). On the other hand, the pooled data from all six isolates showed significant differences in invasion efficiency in trypsin (Fig 2C, *P = 0.0029*) and chymotrypsin (Fig 2D, *P = 0.0009*) treated erythrocytes. Across donors, variations in invasion profiles (defined as the combination of sensitivity to the three enzymes) were mainly driven by sensitivity to trypsin and chymotrypsin treatments, since neuraminidase sensitivity remained unchanged in both individual and pooled parasite invasion rates (Tables A2–A7).

### Relationship between receptor density and sensitivity to enzyme treatment

Having shown that the sensitivity to enzyme treatment varies from donor to donor, we sought to investigate the relationship between the levels of erythrocyte surface receptors and the sensitivity to enzyme treatment. As expected, the expression levels of key erythrocyte surface receptors significantly varied between donors (Fig 3). The median fluorescence intensity (MFI) of labelled antibodies on enzyme-treated erythrocytes was measured and expressed as the percentage of the MFI of the corresponding untreated donor erythrocytes. No significant association was observed between receptor density and the efficiency of enzyme treatment (Fig 4). Given that trypsin and chymotrypsin treatments affect the peptide backbones of the target erythrocyte receptors, we hypothesized that the observed variations in invasion profile after trypsin and chymotrypsin treatments might be driven by the differential expression of receptors on the surface of donor erythrocytes. However, no significant correlation was observed between receptor density and invasion efficiency in enzyme-treated erythrocytes (Fig A1).

**Figure 3:**
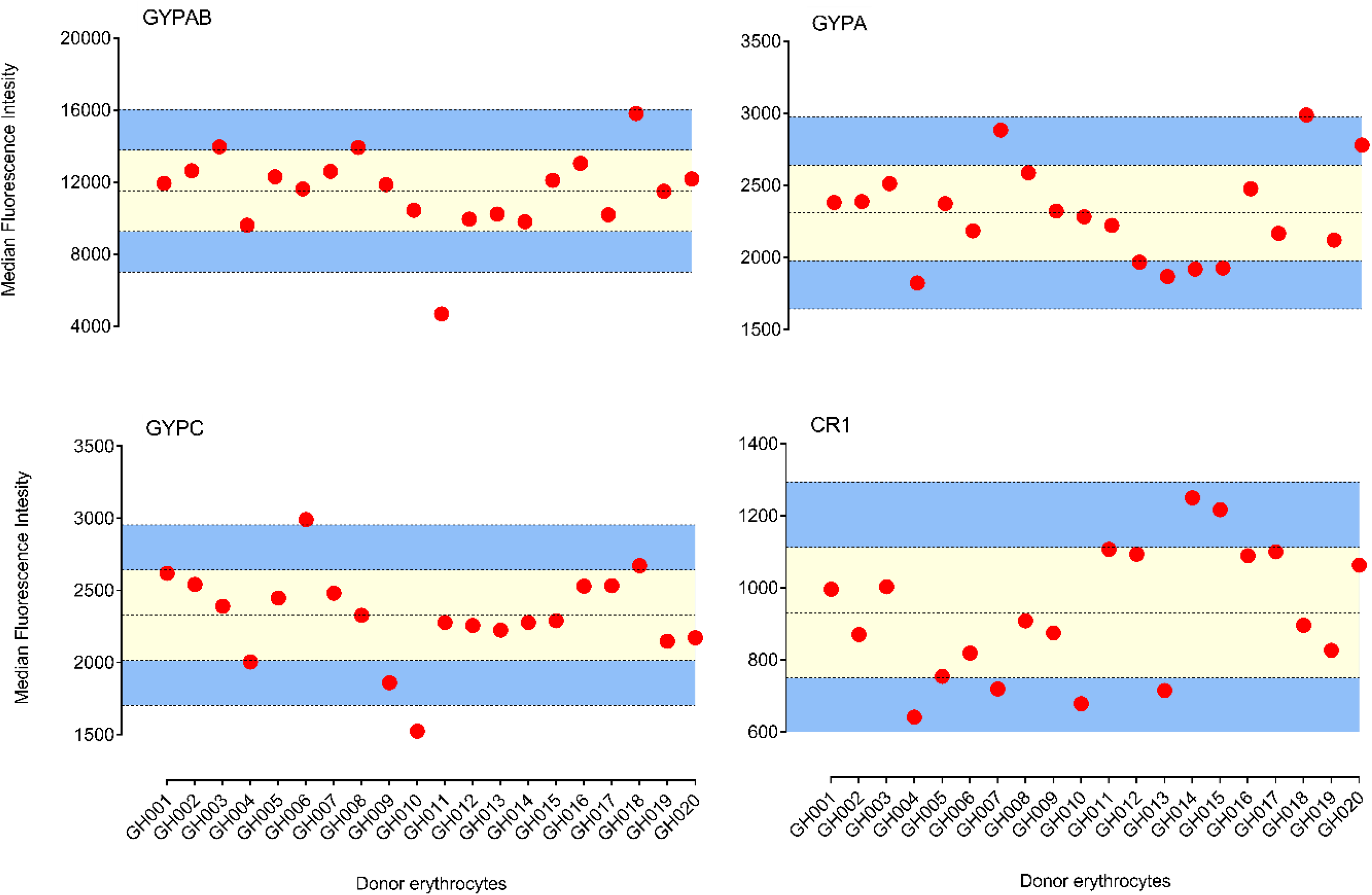
Variation of surface expression of erythrocyte receptors between donors. Dot plots of relative median fluorescent intensity (Y-axis) associated with fluorescently labelled antibodies against specific erythrocyte receptors (GYPAB, GYPA, GYPC and CR1) from 20 blood donors (X-axis). Data were acquired by a BD LSR Fortessa X-20 flow cytometer and processed with FlowJo v.10.01. The data were stratified following normalization and donors were classified as high (Mean + 2SD, upper line), medium (between Mean ± SD, yellow area) and low expressers (Mean-2SD, bottom line).Graphs were plotted using GraphPad Prism v.8.01.

**Figure 4:**
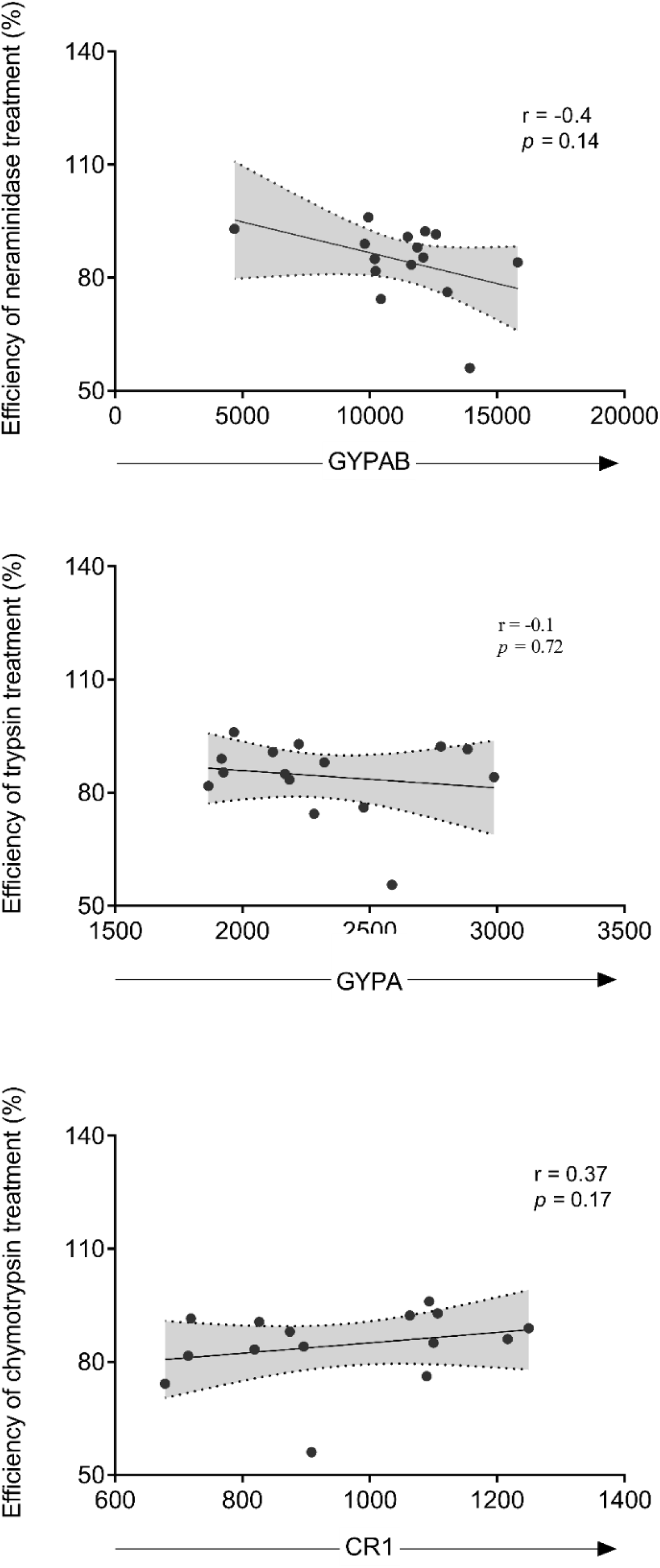
Correlation of erythrocyte receptor densities with the efficiency of enzyme treatment. The density of erythrocyte surface receptors, labelled with specific monoclonal antibodies prior to, and following enzyme treatment was quantified by flow cytometry using fluorescently labelled secondary antibodies. The efficiency of enzyme treatment was then expressed as follows: 100 – (MFI_Treated_ Erythrocytes / MFI_Control_ Erythrocytes × 100). Experiments were performed in triplicates and the resulting data were analysed with FlowJo v.10.01 and the Spearman R correlation were performed using Graph Pad Prism v.8.01.

### Relationship between receptor density and *P. falciparum* invasion efficiency

Given the earlier reports on the relationship between the expression level of specific erythrocyte surface receptors and the invasion efficiency of *P. falciparum* strains (29–33), we sought to rule out any effect of enzyme treatment that could mask a possible correlation between these variables. We selected five different donors (Donors 6-10, refer to Fig 3), with different levels of expression of individual receptors and conducted antibody-mediated invasion inhibitory assays using different concentrations of specific anti-human monoclonal antibodies. As expected, there was a concentration-dependent invasion inhibition by GYPC, CR1 and DAF antibodies. Overall, there were significant differences in the invasion rates across all concentrations (Figs 5A-C). A similar pattern was observed across all antibodies, with the lowest invasion rates always recorded from donor GH006, while donor GH008 recorded the highest invasion rates (Figs 5A-C). However, no linear relationship was observed between the invasion efficiency and the relative abundance of GPC, CR1 or DAF erythrocyte surface receptors (Figs 5A-F), suggesting invasion efficiency may be influenced by other donor-specific cellular properties of erythrocytes.

**Figure 5:**
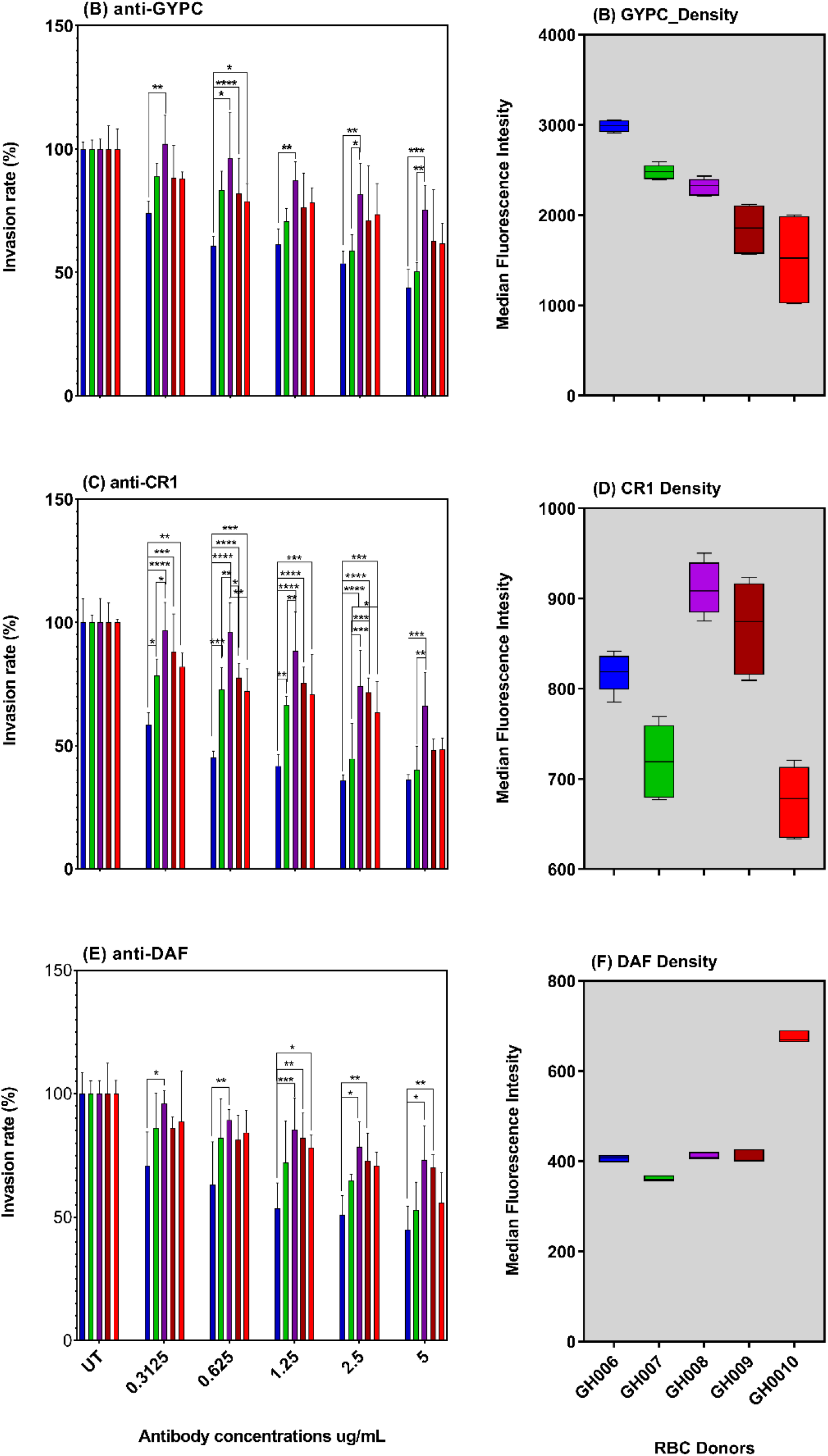
Antibody-dependent invasion inhibition assays in donor erythrocytes with different levels of surface antigens. Schizont-infected erythrocytes were co-incubated with antibody-sensitized uninfected erythrocytes from donors expressing different levels of erythrocyte receptors. The parasites’ DNA was labelled with Hoechst 33342, 18-24 hours’ post-incubation and the resulting parasitemia was quantified by flow cytometry. For each donor, the parasitemia in the corresponding mock-treated erythrocytes was used to ascertain the antibody-dependent invasion inhibitory, following normalization using a single erythrocyte donor (also used for the parasites *in vitro* culturing). A-C represent the antibody-dependent invasion inhibition for two different *P. falciparum* isolates, 3D7 and MISA011. D-F represent the differential expression of erythrocyte receptors assessed in A-C.

### Effects of ABO blood group antigens and hemoglobin genotypes on *P. falciparum* invasion phenotype

Protection against malaria clinical manifestations has for long been associated with erythrocyte surface antigens such as those of the ABO blood group system, as well as erythrocyte hemoglobin defects (18, 19, 34–36). We assessed the relative contribution of donor blood group (Figs 6A, C and E) and hemoglobin genotype (Figs 6B, D and F) in the observed variation in invasion phenotypes. Except for chymotrypsin treatment (P *= 0.0053*), there was no significant difference in the invasion efficiency into erythrocytes of different blood groups (Figs 6A, C and E). This pattern was conserved when comparing invasion efficiencies into erythrocytes of different hemoglobin genotypes, where there was only significant difference following chymotrypsin treatment (*P = 0.0231*) (Figs 6B, D and F). In addition, we conducted a multilinear regression analysis with the donor blood group and hemoglobin genotype as independent variable and showed that none of these variables was a strong predictor of the observed invasion efficiency (data not shown). These findings further suggest that the observed differences are likely due to other intrinsic characteristics specific to each individual donor rather than commonly shared erythrocyte features.

**Figure 6:**
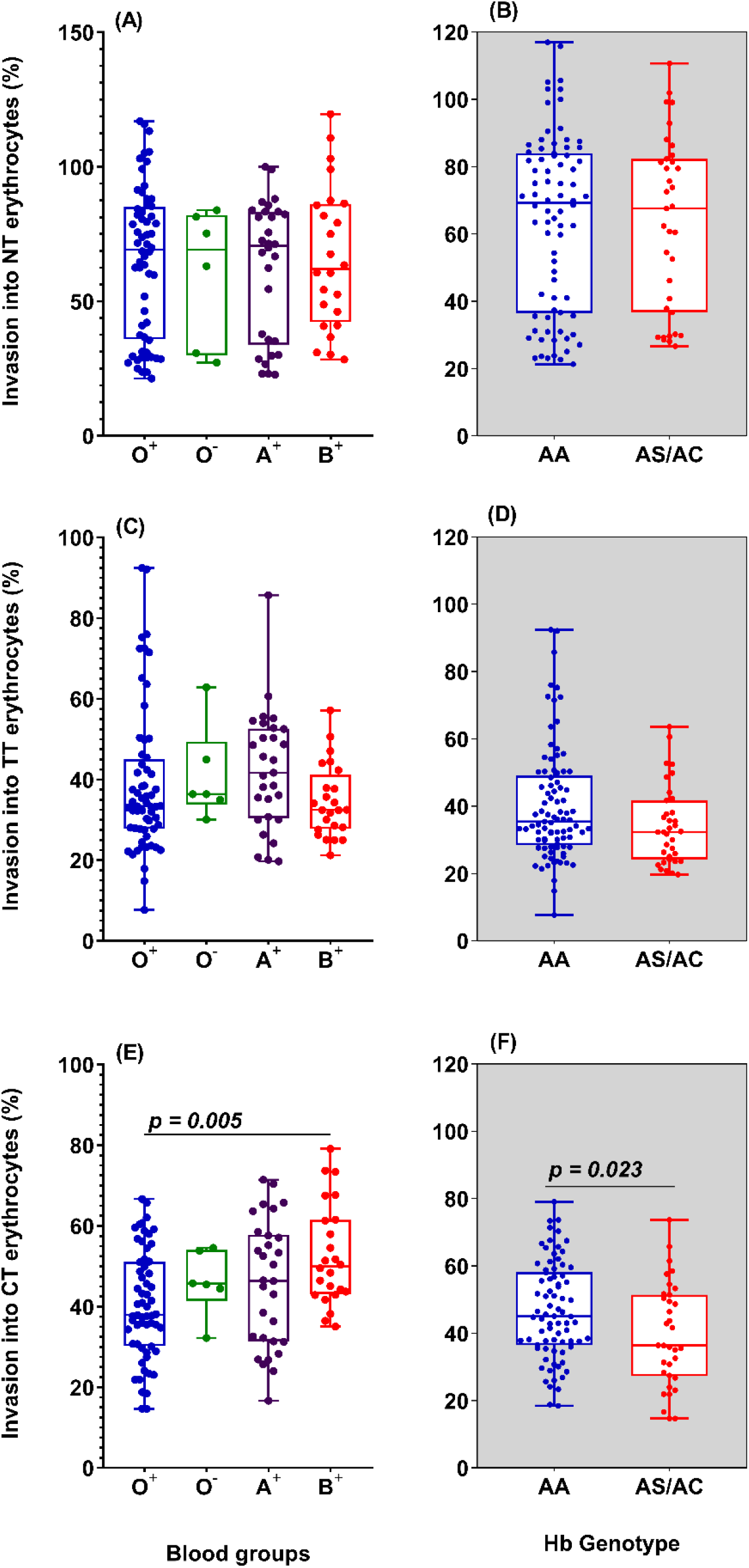
Relationship between blood group or haemoglobin genotype and invasion into enzyme-treated erythrocytes. Erythrocytes from donors of different blood groups (Left panel; O^+^ = 10, O^−^ = 1, A^+^ = 5 and B^+^ = 4) or haemoglobin genotypes (Right panel; AA = 14, AS/AC = 5) were treated with either neuraminidase (A-B), trypsin (C-D) or chymotrypsin (E-F) and coincubated with schizont-infected *P. falciparum* isolates for 18-24h. For each set of donor erythrocytes, the invasion efficiency was assessed using flow cytometry and compared to that of mock-treated control erythrocytes. The data were analysed using Graph Pad Prism v.8.01. Differences in the invasion efficiency into erythrocytes of different blood group were assessed using the Kruskal Wallis test and where significant, pairwise comparisons were performed with the Dunn’s multiple comparison test. The Mann-Whitney test was used to compute for differences in the invasion efficiency into erythrocytes harbouring different haemoglobin genotypes.

## Discussion

This study was undertaken to investigate the effect of blood donor variability in *P. falciparum* IPAs. *P. falciparum* invasion of erythrocytes is a multistep process which includes a tightly regulated interaction between the parasite-derived ligands and specific receptors on the host cell surface. These interactions occur in a functionally redundant manner and therefore define the parasite’s invasion pathway.

Here, we present results from investigations of the effect of key erythrocyte phenotypic attributes in *P. falciparum* IPAs. Using erythrocytes from 20 unrelated donors with different blood groups and hemoglobin genotypes, we showed a wide range of variability in the invasion efficiency of *P. falciparum* strains following enzyme treatment of erythrocytes from distinct donors. We observed a more pronounced variation in invasion profiles, across donors, following treatment with either trypsin or chymotrypsin as compared to neuraminidase. This could partially be as a result of the mode of action of the individual enzymes. Given that trypsin and chymotrypsin are more promiscuous than neuraminidase, they could affect a wider range of receptors, including some with unknown function during the invasion. Besides, this could also be as a result of a small number of binding sites still present following enzyme treatment. As we found that cells from individual donors presented different levels of sensitivity to enzyme treatment, we postulated that erythrocyte receptors are differentially expressed on the surface of individual donor cells, thus driving the sensitivity to enzyme treatment. Although the expression level of erythrocyte receptors varied across donors, we observed no relationship between receptor density and sensitivity to enzyme treatment. Knockdown experiments of both basigin and CR1 have previously shown a linear relationship between receptor density and invasion efficiency into erythrocytes (29, 30). Here, there is a possibility that the absence of a relationship between receptor density and invasion efficiency was due to the effect of enzyme treatment of the erythrocytes. To rule out any effect of enzyme treatment, erythrocytes from five different donors were used in an antibody-dependent invasion inhibition assay without any prior enzyme treatment. Increasing concentrations of antibodies limits the number of receptors available for parasite invasion, therefore mimicking the effect of receptor knockdown at the individual level. In the case of a positive linear relationship between these two variables, one would expect higher invasion rates across all dilutions in individuals with a higher expression of a given receptor. Although there was an evident dose-dependent invasion inhibition by the various antibodies when donors were taken individually, there was no apparent linear relationship between receptor density and invasion efficiency. This suggests that receptor density is not the only factor influencing invasion efficiency in erythrocytes of different origin.

The relationship between *P. falciparum* and the ABO blood group system or hemoglobin genotypes remains a fascinating subject for researchers. Despite the small number of donors included in this study, we sought to investigate a possible relation between the observed invasion phenotype and the donor blood group or hemoglobin genotype. Overall, across all donors, invasion into erythrocytes of different blood groups or hemoglobin genotypes only differed significantly following chymotrypsin treatment. While majority of the hemoglobin variants conferring protection against malaria have been shown not to significantly affect erythrocyte invasion *in vitro* (37), several studies have reported the parasite preference for O^+^ blood group *in vitro* (38, 39). More recently, different *P. falciparum* strains have been shown to differentially invade erythrocytes of the same blood group from thirty different donors (40), suggesting that variation in invasion efficiency could be driven by other host-specific features that are yet to be defined.

Understanding the interplay between merozoite attachment and the changes in the erythrocyte biophysical properties would add complementary information about the host cell contribution to invasion. Using a mechanistic model for parasite-membrane adhesion, Hillringhaus and colleagues showed that the adhesion of the invading merozoite is further strengthened by erythrocyte deformability upon primary contact (41). This agrees with earlier studies suggesting the occurrence of biophysical changes in the host cell upon primary contact with external ligands (42–44). This suggests a more or less active contribution of the host erythrocyte to the invasion efficiency, through activation of downstream signaling pathways following the primary contact with the invading merozoite. Together, these findings emphasize the need for more robust assays to characterize the actual contribution of the host cell during *P. falciparum* invasion. Finally, given the heterogeneity of peripheral erythrocytes (45), and the consequent changes in the expression of different erythrocyte surface markers (46, 47), it is of utmost importance to consider the relative contribution of the different erythrocyte sub-populations circulating in the peripheral blood.

## Author Contributions

LGT, YA and GAA conceived the study; LGT and PBN performed the experiments LGT, PBN, YA and GAA analyzed the data and drafted the manuscript; YA, GAA, MN and KAK supervised the study. All authors critically reviewed and edited the manuscript.

## Acknowledgements

This work is part of the assay standardization efforts of the West African merozoite invasion network (WAMIN) consortium, and we are grateful to members for contributing ideas to this work. We also thank Dr Manoj T. Duraisingh, Dr Estela Shabani and Dr Martha Clark of the Harvard School of Public Health for the insightful discussions during the study design.

## Supplemery information

## Appendixes

**Figure A1:**
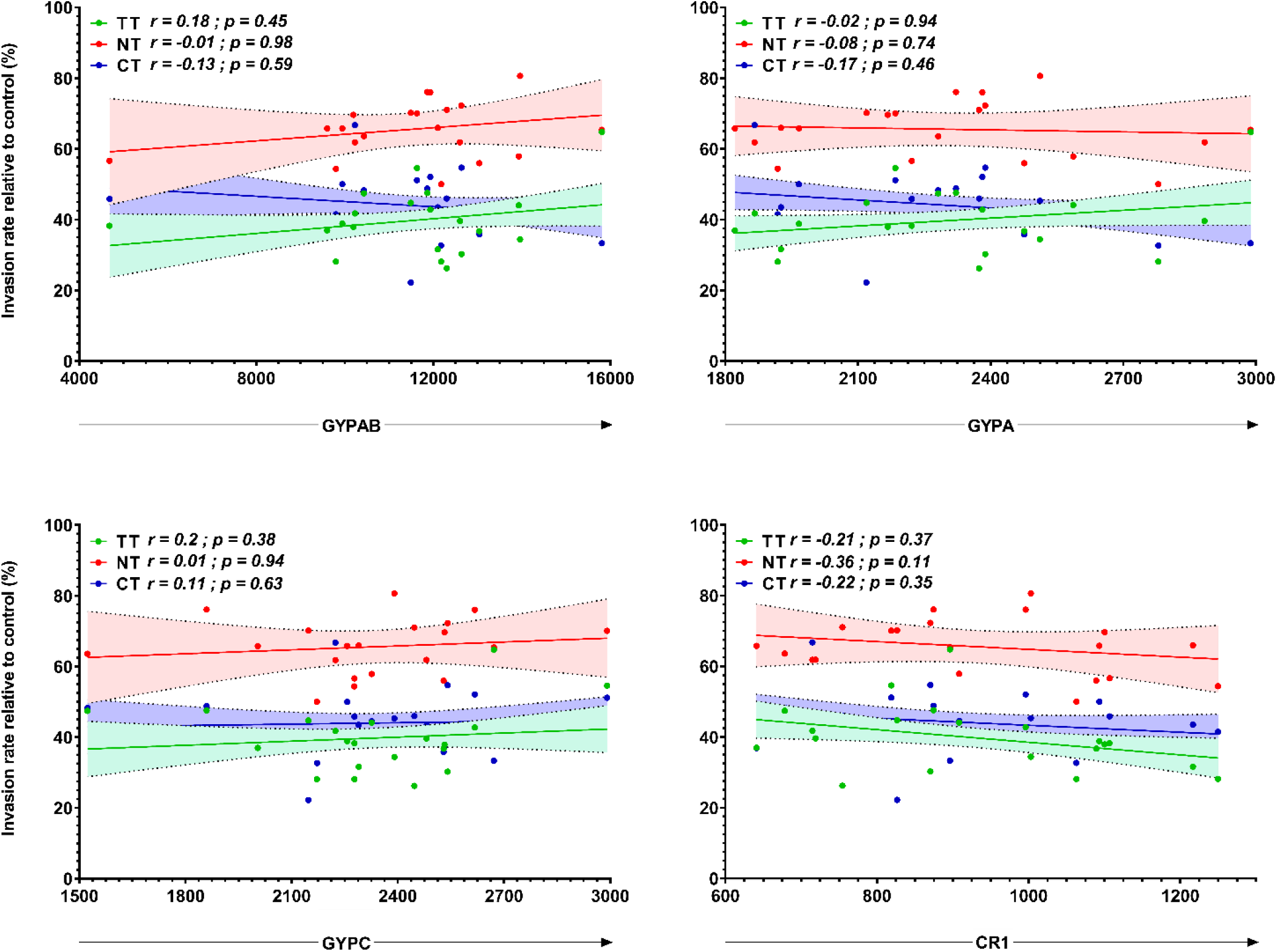
Correlation of receptor density with invasion efficiency into enzyme-treated erythrocytes. The Spearman correlation test was used to assess the relationship between the density of individual erythrocyte surface receptors (X-axis) and the invasion efficiency following enzyme treatment of the same erythrocyte (Y-axis). The data were acquired as MFI for the receptor density and per cent parasitemia relative to untreated control erythrocytes for the invasion efficiency and the graphs were plotted using Graph Pad Prism v.8.01.

**Table A1:** Clinical and hematological indices of study participants

| ID | Gender | Blood group | Sickling | Hb genotype | G6PD | WBC (10<sup>9</sup>/L) | RBC (10<sup>12</sup>/L) | HGB (g/dL) | HCT (%) | MCV (fL) | MCH (pg) | MCHC (g/dL) | PLT (10<sup>9</sup>/L) |
| --- | --- | --- | --- | --- | --- | --- | --- | --- | --- | --- | --- | --- | --- |
| **GH000\*\*** | M | O+ | - | AA | Normal | 4.34 | 5.16 | 13.0 | 42.8 | 83.0 | 25.2 | 30.4 | 153 |
| **GH001** | F | O+ | - | AA | Normal | 7.98 | 3.93 | 11.5 | 38.9 | 98.9 | 29.3 | 29.6 | 342 |
| **GH002** | M | B+ | - | AA | Normal | 5.91 | 6.04 | 15.0 | 51.5 | 85.2 | 24.8 | 29.1 | 231 |
| **GH003** | M | B+ | + | AS | Normal | 3.16 | 5.34 | 15.1 | 46.7 | 87.4 | 28.3 | 32.3 | 220 |
| **GH004** | M | O+ | + | AS | Normal | 4.38 | 6.26 | 13.7 | 44.8 | 71.5 | 21.9 | 30.6 | 218 |
| **GH005** | M | O+ | - | AA | Normal | 6.82 | 5.31 | 14.3 | 47.1 | 88.7 | 26.9 | 30.4 | 202 |
| **GH006** | F | A+ | - | AA | Normal | 6.88 | 3.83 | 12.3 | 39.4 | 102.8 | 32.1 | 31.2 | 273 |
| **GH007** | M | O- | - | AA | Normal | 5,07 | 4,77 | 14,5 | 43,6 | 91,4 | 30,4 | 33,3 | 228 |
| **GH008** | M | A+ | - | AA | NA | NA | NA | NA | NA | NA | NA | NA | NA |
| **GH009** | M | O+ | - | AA | Normal | 5,59 | 3,76 | 13,3 | 39,9 | 106,2 | 35,4 | 33,3 | 317 |
| **GH010** | M | A+ | + | AS | Normal | 5.57 | 6.30 | 16.3 | 51.7 | 82.0 | 25.9 | 31.5 | 258 |
| **GH011** | M | O+ | - | AA | Normal | 3.86 | 5.21 | 15.4 | 48.3 | 92.7 | 29.6 | 31.9 | 169 |
| **GH012** | M | O+ | - | AA | Normal | 4.74 | 5.94 | 15.2 | 49.5 | 83.4 | 25.6 | 30.7 | 197 |
| **GH013** | F | B+ | - | AA | Normal | 5.53 | 4.57 | 13.5 | 43.0 | 94.0 | 29.5 | 31.4 | 244 |
| **GH014** | F | B+ | + | AS | Normal | 5.73 | 4.61 | 12.9 | 40.2 | 87.1 | 28.0 | 32.1 | 251 |
| **GH015** | M | O+ | - | AA | Normal | 7.90 | 4.95 | 14.6 | 46.8 | 94.6 | 29.5 | 31.2 | 253 |
| **GH016** | M | A+ | - | AC | Normal | 5.49 | 4.74 | 13.2 | 39.6 | 83.5 | 27.8 | 33.3 | 244 |
| **GH017** | M | A+ | - | AA | Full Defect | 3,02 | 3,85 | 9,9 | 31,5 | 81,8 | 25,7 | 31,4 | 209 |
| **GH018** | F | O+ | NA | NA | NA | NA | NA | NA | NA | NA | NA | NA | NA |
| **GH019** | M | O+ | - | AC | Normal | 7.70 | 5.48 | 14.6 | 45.2 | 82.5 | 26.6 | 32.3 | 260 |
| **GH020** | M | O+ | - | AA | Normal | 5.94 | 5.35 | 15.7 | 49.1 | 91.8 | 29.3 | 32.0 | 187 |

\*\* Erythrocytes from this donor were used for routine parasite culturing and for data normalization

**Table A2:** MISA010. Summary of invasion profiles of individual *P. falciparum* isolates into enzyme-treated erythrocyte from different donors. Data represent mean values ± the standard deviation from two independent experiments conducted in triplicates. A cut-off of 50% was taken as a threshold to classify the observed profile as sensitive (≤50%) or resistance (>50%) for each given treatment.

| % invasion relative to untreated control |  |  |  |  |  |
| --- | --- | --- | --- | --- | --- |
| Donor Erythrocyte | Gender | Invasion profile | Neuraminidase (250 mU/ml) | Trypsin (1mg/ml) | Chymotrypsin (1mg/ml) |
| O+/AA | F | NrTsCr | 103.1 ± 16.3 | 41.7 ± 4.1 | 60.4 ± 17.0 |
| B+/AA | M | NrTsCr | 75.0 ± 8.1 | 25.0 ± 7.0 | 61.3 ± 11.2 |
| B+AS | M | NrTsCs | 119.6 ± 22.4 | 28.6 ± 4.2 | 42.9 ± 8.6 |
| O+AS | M | NrTsCs | 62.5 ± 6.1 | 35.4 ± 15.6 | 35.4 ± 18.9 |
| O+AA | M | NrTsCr | 75.7 ± 23.3 | 23.6 ± 4.3 | 51.7 ± 19.5 |
| A+/AA | F | NrTrCr | 71.43 ± 18.31 | 85.71 ± 1,23 | 57.14 ± 17.73 |
| O-/AA | M | NrTsCs | 75.22 ± 9.87 | 44.95 ± 1.47 | 45.45 ± 12.55 |
| A+/AA | M | NrTrCs | 66.66 ± 17.36 | 55.55 ± 4.78 | 46.29 ± 4,78 |
| O+/AA | M | NrTrCr | 80.55 ± 13.58 | 58.33 ± 29.48 | 65.74 ± 14.3 |
| A+/AS | M | NrTrCr | 81.31 ± 5.62 | 52.52 ± 16.52 | 57.57 ± 16.26 |
| O+/AA | M | NrTrCr | 81.62 ± 5,87 | 72.46 ± 14.54 | 51.0 ± 5.85 |
| O+/AA | M | NrTrCs | 75.12 ± 2.17 | 71.52 ± 13.18 | 40.59 ± 7.25 |
| B+/AA | M | NrTsCr | 54.35 ± 4.24 | 47.06 ± 8.5 | 67.64 ± 7.15 |
| B+/AS | F | NrTsCs | 60.71 ± 0.92 | 25 ± 13.04 | 41.67 ± 16.34 |
| O+/AA | M | NrTsCs | 71.23 ± 21.05 | 32 ± 15.31 | 36 ± 7.37 |
| A+/AC | M | NrTsCr | 83.33 ± 11.72 | 41.67 ± 13.44 | 50.42 ± 13.1 |
| A+/AA | M | NrTrCr | 83.33 ± 14.91 | 50.34 ± 19.72 | 52.5 ± 5.1 |
| O+/AA | F | NrTrCs | 85.36 ± 16.71 | 65.13 ± 12.19 | 45.65 ± 5.59 |
| O+/AC | M | NrTsCs | 79.48 ± 11.14 | 35.90 ± 14.56 | 21.88 ± 9.97 |
| O+/AA | M | NrTsCs | 63.49 ± 15.69 | 22.22 ± 0.65 | 34.75 ± 7.70 |

**Table A3:** MISA011. Summary of invasion profiles of individual *P. falciparum* isolates into enzyme-treated erythrocyte from different donors. Data represent mean values ± the standard deviation from two independent experiments conducted in triplicates. A cut-off of 50% was taken as a threshold to classify the observed profile as sensitive (≤50%) or resistance (>50%) for each given treatment.

| **Donor**<br>**erythrocytes** | **Gender** | **% invasion relative to untreated control** |  |  |  |
| --- | --- | --- | --- | --- | --- |
|  |  | **Invasion**<br>**profile** | **Neuraminidase**<br>**(250 mU/ml)** | **Trypsin**<br>**(1mg/ml)** | **Chymotrypsin**<br>**(1mg/ml)** |
| **O+/AA** | F | NrTsCr | 115.89 ± 2.33 | 42.39 ± 10.26 | 56.75 ± 26.27 |
| **B+/AA** | M | NrTsCr | 103.03 ± 8.11 | 25.06 ± 9.42 | 50.35 ± 6.36 |
| **B+/AS** | M | NrTsCs | 110.71 ± 24.1 | 35.71 ± 19.19 | 46.43 ± 17.56 |
| **O+/AS** | M | NrTsCs | 105.14 ± 8.52 | 30.28 ± 8.39 | 40 ± 14.97 |
| **O+/AA** | M | NrTsCs | 101.96 ± 18.13 | 33.44 ± 5.92 | 35.89 ± 9.95 |
| **A+/AA** | F | NrTrCs | 99.06 ± 14.35 | 55.14 ± 15.69 | 46.48 ± 7.23 |
| **O-/AA** | M | NrTsCs | 83.89 ± 14.81 | 30.09 ± 11.83 | 32.2 ± 13.89 |
| **A+/AA** | M | NrTsCs | 72.55 ± 11.24 | 32.84 ± 8.9 | 42.53 ± 6.85 |
| **O+/AA** | M | NrTsCs | 105.60 ± 9.64 | 37.58 ± 9.36 | 37.36 ± 9.78 |
| **A+/AS** | M | NrTsCs | 88.04 ± 10.32 | 36.14 ± 21.78 | 31.26 ± 11.93 |
| **O+/AA** | M | NrTsCs | 59.76 ± 6.28 | 23.15 ± 6.24 | 29.67 ± 5.31 |
| **O+/AA** | M | NrTsCs | 71.73 ± 9.29 | 33.30 ± 11.85 | 43.01 ± 16.62 |
| **B+/AA** | M | NrTrCs | 81.81 ± 3.84 | 50.65 ± 9.00 | 79.11 ± 7.65 |
| **B+/AS** | F | NrTsCs | 60.53 ± 3.84 | 26.29 ± 9.55 | 36.45 ± 13.45 |
| **O+/AA** | M | NrTrCs | 91.38 ± 16.01 | 29.79 ± 14.67 | 37.70 ± 11.05 |
| **A+/AC** | M | NrTsCs | 72.56 ± 3.98 | 20.76 ± 2.02 | 28.31 ± 1.70 |
| **A+/AA** | M | NrTsCs | 82.31 ± 7.00 | 19.72 ± 3.85 | 32.60 ± 4.98 |
| **O+/AA** | F | NrTrCs | 83.20 ± 13.44 | 72.56 ± 7.87 | 26.05 ± 2.31 |
| **O+/AC** | M | NrTsCs | 113.26 ± 12.97 | 32.33 ± 19.87 | 27.45 ± 12.71 |
| **O+/AA** | M | NrTsCs | 66.80 ± 3.25 | 23.33 ± 7.43 | 34.31 ± 11.03 |

**Table A4:** MISA018. Summary of invasion profiles of individual *P. falciparum* isolates into enzyme-treated erythrocyte from different donors. Data represent mean values ± the standard deviation from two independent experiments conducted in triplicates. A cut-off of 50% was taken as a threshold to classify the observed profile as sensitive (≤50%) or resistance (>50%) for each given treatment.

| Donor<br>Erythrocyte | Gender | % invasion relative to untreated control |  |  |  |
| --- | --- | --- | --- | --- | --- |
|  |  | Invasion<br>profile | Neuraminidase<br>(250 mU/ml) | Trypsin<br>(1mg/ml) | Chymotrypsin<br>(1mg/ml) |
| **O+/AA** | F | NrTsCr | 90.48 ± 7.84 | 38.28 ± 5.14 | 44.8 ± 5.13 |
| **B+/AA** | M | NrTsCr | 87.47 ± 11.81 | 27.58 ± 12.07 | 38.24 ± 12.42 |
| **B+AS** | M | NrTsCs | 99.08 ± 6.46 | 34.31 ± 14.28 | 43.72 ± 11.36 |
| **O+AS** | M | NrTsCs | 78.91 ± 4.89 | 33.15 ± 13.31 | 35.51 ± 6.58 |
| **O+AA** | M | NrTsCr | 92.89 ± 4.59 | 27.46 ± 6.22 | 35,45 ± 5.68 |
| **A+/AA** | F | NrTrCr | 75.59 ± 4.59 | 66.67 ± 7.07 | 65.38 ± 16.79 |
| **O-/AA** | M | NrTsCs | 63.09 ± 11.75 | 36.36 ± 8.76 | 45.80 ± 9.87 |
| **A+/AA** | M | NrTrCr | 71.21 ± 18.51 | 53.98 ± 16.55 | 55.21 ± 9.61 |
| **O+/AA** | M | NrTrCr | 116.98 ± 24.10 | 75.26 ± 30.78 | 66.67 ± 9,24 |
| **A+/AS** | M | NrTrCr | 81.34 ± 25.29 | 52.76 ± 26.97 | 58.5 ± 13.99 |
| **O+/AA** | M | NrTsCs | 64.61 ± 7.60 | 46.15 ± 2.65 | 43.25 ± 10.92 |
| **O+/AA** | M | NrTrCr | 84.31 ± 14.20 | 50.15 ± 15.49 | 60.74 ± 8.94 |
| **B+/AA** | M | NrTrCs | 85.71 ± 12.65 | 57.14 ± 4.92 | 42.86 ± 5,53 |
| **B+/AS** | F | NrTsCr | 52.57 ± 10.63 | 21.2 ± 2.99 | 54.55 ± 12.96 |
| **O+/AA** | M | NrTsCr | 68.36 ± 27.85 | 34.48 ± 8,51 | 62.07 ± 16.79 |
| **A+/AC** | M | NrTsCs | 54.54 ± 5,63 | 20.09 ± 4,30 | 36.36 ± 14.76 |
| **A+/AA** | M | NrTsCr | 85.71 ± 2.95 | 45.71 ± 6.76 | 71.42 ± 17.21 |
| **O+/AA** | F | NrTrCs | 62.50 ± 15.86 | 50.36 ± 16.45 | 40.93 ± 23.05 |
| **O+/AC** | M | NrTsCs | 73.85 ± 10.27 | 32.31 ± 11.19 | 23.08 ± 7.96 |
| **O+/AA** | M | NrTsCs | 60.29 ± 5.31 | 27.94 ± 9.33 | 37.65 ± 10.11 |

**Table A5:** 3D7. Summary of invasion profiles of individual *P. falciparum* isolates into enzyme-treated erythrocyte from different donors. Data represent mean values ± the standard deviation from two independent experiments conducted in triplicates. A cut-off of 50% was taken as a threshold to classify the observed profile as sensitive (≤50%) or resistance (>50%) for each given treatment.

| % invasion relative to untreated control |  |  |  |  |  |
| --- | --- | --- | --- | --- | --- |
| Donor Erythrocyte | Gender | Invasion profile | Neuraminidase (250 mU/ml) | Trypsin (1mg/ml) | Chymotrypsin (1mg/ml) |
| O+/AA | F | NrTsCs | 86.54 ± 9.82 | 43.76 ± 11.41 | 49.9 ± 7.22 |
| B+/AA | M | NrTsCs | 79.24 ± 16.54 | 30.05 ± 9.74 | 44.26 ± 4.43 |
| B+/AS | M | NrTsCs | 86.36 ± 11.64 | 31.76 ± 8.91 | 48.37 ± 7.05 |
| O+/AS | M | NrTsCs | 88.02 ± 15.45 | 30.57 ± 2.00 | 47.19 ± 4.61 |
| O+/AA | M | NrTsCs | 88.07 ± 14.44 | 23.25 ± 11.51 | 48.70 ± 7.05 |
| A+/AA | F | NrTsCs | 86.93 ± 8.26 | 48.52 ± 7.97 | 45 ± 8.21 |
| O-/AA | M | NrTsCs | 81.39 ± 8.92 | 35 ± 8.21 | 44.44 ± 6.34 |
| A+/AA | M | NrTsCs | 69.98 ± 6.48 | 40.98 ± 5.75 | 43.06 ± 3.06 |
| O+/AA | M | NrTsCs | 78.67 ± 6.69 | 34.17 ± 6.44 | 36.61 ± 7.87 |
| A+/AS | M | NrTsCs | 62.36 ± 9.78 | 30.00 ± 6.27 | 31.37 ± 5.95 |
| O+/AA | M | NrTsCr | 68.61 ± 8.55 | 28.72 ± 9.3 | 59.18 ± 7.89 |
| O+/AA | M | NrTsCr | 74.68 ± 9.55 | 33.03 ± 6.81 | 58.81 ± 6.34 |
| B+/AA | M | NrTsCs | 63.43 ± 43 | 32.37 ± 6.19 | 45.09 ± 8.43 |
| B+/AS | F | NrTsCr | 67.57 ± 10.84 | 28.14 ± 21.94 | 57.95 ± 18.05 |
| O+/AA | M | NrTsCr | 82.13 ± 11.59 | 27.8 ± 9.09 | 54.52 ± 13.83 |
| A+/AC | M | NrTsCs | 68.17 ± 3.87 | 24.18 ± 2.97 | 24.01 ± 2.04 |
| A+/AA | M | NrTsCs | 100 ± 6.70 | 26.31 ± 0.39 | 25.66 ± 4.72 |
| O+/AA | F | NrTsCs | 69.89 ± 10.98 | 76 ± 15.4 | 24.1 ± 1.04 |
| O+/AC | M | NrTsCs | 99.23 ± 3.79 | 36.71 ± 4.09 | 14.68 ± 7.78 |
| O+/AA | M | NrTsCs | 51.87 ± 14.31 | 7.67 ± 3.21 | 30.70 ± 2.69 |

**Table A6:** Dd2. Summary of invasion profiles of individual *P. falciparum* isolates into enzyme-treated erythrocyte from different donors. Data represent mean values ± the standard deviation from two independent experiments conducted in triplicates. A cut-off of 50% was taken as a threshold to classify the observed profile as sensitive (≤50%) or resistance (>50%) for each given treatment.

| % invasion relative to untreated control |  |  |  |  |  |
| --- | --- | --- | --- | --- | --- |
| Donor Erythrocyte | Gender | Invasion profile | Neuraminidase (250 mU/ml) | Trypsin (1mg/ml) | Chymotrypsin (1mg/ml) |
| O+/AA | F | NsTsCs | 30.94 ± 4.34 | 32.25 ± 1.85 | 37.87 ± 4.47 |
| B+/AA | M | NsTsCr | 31.03 ± 3.08 | 37.93 ± 16.82 | 51.72 ± 2.05 |
| B+/AS | M | NsTsCr | 36.71 ± 13.75 | 44.05 ± 5.69 | 51.39 ± 2.05 |
| O+/AS | M | NsTsCr | 31.03 ± 3.08 | 37.93 ± 6.04 | 51.72 ± 2.05 |
| O+/AA | M | NsTsCr | 36.71 ± 5.76 | 44.05 ± 5.69 | 51.4 ± 6.69 |
| A+/AA | F | NsTsCs | 23.08 ± 6.89 | 30.77 ± 6.95 | 38.46 ± 7.94 |
| O-/AA | M | NsTsCr | 27.27 ± 11,56 | 36.06 ± 8.21 | 54.26 ± 4.69 |
| A+/AA | M | NsTsCr | 23.07 ± 13.24 | 38.46 ± 7.94 | 53.85 ± 7.94 |
| O+/AA | M | NsTsCs | 23.80 ± 3.25 | 21.42 ± 5.61 | 28.57 ± 7.38 |
| A+/AS | M | NsTsCr | 26.67 ± 8.27 | 35.53 ± 6.49 | 53.33 ± 5.96 |
| O+/AA | M | NsTsCs | 36.65 ± 2.82 | 17.79 ± 10.54 | 18.43 ± 5.31 |
| O+/AA | M | NsTsCr | 35.57 ± 1.50 | 28.07 ± 3.51 | 56.14 ± 13.54 |
| B+/AA | M | NsTsCr | 41.03 ± 4.13 | 32.5 ± 3.09 | 67.5 ± 18.43 |
| B+/AS | F | NsTsCr | 46.15 ± 2.16 | 42.31 ± 15,75 | 61.54 ± 10.14 |
| O+/AA | M | NsTsCr | 41.03 ± 5.05 | 14.80 ± 6.39 | 59.59 ± 10.87 |
| A+/AC | M | NsTsCs | 28.57 ± 2.83 | 38.09 ± 12.13 | 16.67 ± 2.56 |
| A+/AA | M | NsTsCr | 35.18 ± 11.04 | 35.12 ± 12.59 | 70.37 ± 13.18 |
| O+/AA | F | NsTrCs | 42.10 ± 4.24 | 92.10 ± 4.08 | 23.39 ± 3.51 |
| O+/AC | M | NsTsCs | 29.30 ± 4.96 | 49.82 ± 11.23 | 14.65 ± 4.34 |
| O+/AA | M | NsTsCs | 25.05 ± 11.88 | 24.46 ± 4.59 | 28.59 ± 8.27 |

**Table A7:** W2mef. Summary of invasion profiles of individual *P. falciparum* isolates into enzyme-treated erythrocyte from different donors. Data represent mean values ± the standard deviation from two independent experiments conducted in triplicates. A cut-off of 50% was taken as a threshold to classify the observed profile as sensitive (≤50%) or resistance (>50%) for each given treatment.

| Donor<br>Erythrocyte | Gender | % invasion relative to untreated control |  |  |  |
| --- | --- | --- | --- | --- | --- |
|  |  | Invasion<br>profile | Neuraminidase<br>(250 mU/ml) | Trypsin<br>(1mg/ml) | Chymotrypsin<br>(1mg/ml) |
| **O+/AA** | F | NsTsCr | 29.14 ± 3.37 | 36.05 ± 11.15 | 58.02 ± 14.32 |
| **B+/AA** | M | NsTsCr | 28.36 ± 6.49 | 34.14 ± 9.6 | 73.41 ± 10.43 |
| **B+AS** | M | NsTsCs | 30.16 ± 5.17 | 37.67 ± 8.84 | 49.53 ± 12.87 |
| **O+AS** | M | NsTsCs | 28.89 ± 3.61 | 38.52 ± 0 | 48.15 ± 3.1 |
| **O+AA** | M | NsTsCs | 29.63 ± 3.13 | 25.92 ± 3.89 | 44.44 ± 2.88 |
| **A+/AA** | F | NsTrCr | 35.71 ± 8.21 | 50.32 ± 3,69 | 64.28 ± 13.29 |
| **O-/AA** | M | NsTrCr | 30.77 ± 9.54 | 62.82 ± 14.15 | 53.85 ± 4.42 |
| **A+/AA** | M | NsTrCr | 22.72 ± 4.06 | 54.54 ± 0.78 | 63.64 ± 17.84 |
| **O+/AA** | M | NsTsCr | 23.67 ± 6.49 | 45.45 ± 7,33 | 51.14 ± 10.58 |
| **A+/AS** | M | NsTrCr | 37.87 ± 3.91 | 60.61 ± 11.37 | 65.75 ± 22.12 |
| **O+/AA** | M | NsTsCs | 27.08 ± 13.13 | 22.32 ± 12.01 | 18.75 ± 17.40 |
| **O+/AA** | M | NsTsCr | 37.44 ± 10.42 | 27.78 ± 2.14 | 55.56 ± 17.46 |
| **B+/AA** | M | NsTsCr | 48.82 ± 7.63 | 44.44 ± 7.63 | 73.73 ± 18.10 |
| **B+/AS** | F | NsTsCs | 40.79 ± 14.31 | 32.46 ± 2.72 | 35.05 ± 16.70 |
| **O+/AA** | M | NsTsCs | 31.25 ± 17.39 | 37.5 ± 14.82 | 41.44 ± 15.00 |
| **A+/AC** | M | NsTsCs | 29.83 ± 11.68 | 48.72 ± 8.18 | 26.76 ± 4.03 |
| **A+/AA** | M | NsTsCs | 30.08 ± 9.89 | 44.84 ± 3.37 | 26.89 ± 3.55 |
| **O+/AA** | F | NsTrCs | 46.38 ± 3.56 | 92.44 ± 7.85 | 37.61 ± 4.31 |
| **O+/AC** | M | NsTrCs | 28.07 ± 4.01 | 63.64 ± 8.73 | 21.832 ± 5.38 |
| **O+/AA** | M | NsTsCs | 21.33 ± 0.49 | 41.33 ± 8.39 | 30.11 ± 4.87 |

**Key:** N: neuraminidase, T: trypsin, C: chymotrypsin, s: sensitive, r: resistant Data represent mean values ± the standard deviation from two independent experiments conducted in triplicates. A cut-off of 50% was taken as a threshold to classify the observed profile as sensitive (≤50%) or resistance (>50%) for each given treatment.

